# Bacteria orchestrate gametophyte growth, oogenesis and sporophyte development in *Saccharina latissima* in a sex-dependent manner

**DOI:** 10.64898/2026.04.17.718847

**Authors:** Otto P. van der Linden, Peter A.C. van Gisbergen, Dylan Selles, Detmer Sipkema, Tijs Ketelaar

## Abstract

- Marine organisms, including green and brown macroalgae, exhibit a broad dependency on their microbiome which has been demonstrated in model species including *Ulva compressa* and *Ectocarpus siliculosus* with relatively simple building plans. However, it remains elusive if and how *Saccharina latissima*, a complex brown macroalgae with high degrees of organ and tissue differentiation, is controlled by its microbiome.
- We monitored gametophyte cultures of mixed sexes, induced oogenesis and followed sporophyte development both under axenic conditions and in cultures complemented with bacterial isolates from the sugar kelp core microbiome.
- Female gametophytes generally performed better in the presence of bacteria while males performed worse. Some bacterial isolates inhibit oogenesis in females entirely, whereas others have a stimulating effect. Under axenic conditions sporophytes did form, but growth, pigmentation and the establishment of an apical-basal polarization axis were severely disrupted. These defects could be resolved by complementation with many bacteria from the S. latissima core microbiome.
- Sugar kelp depends heavily on specific bacterial symbionts for growth, reproduction and development and their effect is sex-dependent in gametophytes. This work provides a platform to investigate the precise methods of cross-kingdom communication which has a large potential in the kelp production industry.

## Introduction

Algae represent an evolutionarily diverse group of photosynthetic organisms that occur in marine, fresh water and even terrestrial environments. Three groups of algae, the red, green and brown seaweeds, have acquired multicellularity and are commonly referred to as macroalgae. In all studied seaweeds, bacteria play widely diverse, crucial roles in growth and development. They can inflict disease, affect nutrient availability or improve spore release and settlement, whereas others provide essential growth-promoting factors (Li et al., 2023; Park, 2018; Park et al., 2023; Peng & Li, 2013; Saha & Weinberger, 2019; Wichard, 2015).

For the green seaweed *Ulva compressa*, groundbreaking work has shown that chemicals secreted by diverse groups of bacteria are essential to drive morphogenesis and growth (Egan et al., 2013; Singh & Reddy, 2014). For the brown seaweed, *Ectocarpus siliculosus*, a similar dependency was demonstrated (Taipa et al., 2016). From a wide range of studies on diverse seaweeds, a picture emerges that specific groups of bacteria associate with diverse seaweeds and form a core microbiome, suggesting a conserved key role of microbial partners in driving growth and development of seaweeds (Egan et al., 2013; Li et al., 2023; Singh & Reddy, 2014).

For the brown seaweeds, *Ectocarpus siliculosus* is an emerging model system due to its small size, simple building plan and short life cycle. It lacks, however, the complexity of kelps, larger brown seaweeds that exhibit organ and tissue differentiation, which is remarkably similar to that observed in seed plants. If and how morphogenetic bacteria contribute to the growth and differentiation of large brown seaweeds is still poorly understood. In this work we focus on the interplay between microbiota and the upcoming seaweed crop sugar kelp (*Saccharina latissima*)(Jaugeon et al., 2025).

Because of the high-value biobased products and ecosystem services it can provide, sugar kelp is one of the seaweeds of choice for cultivation in North West European waters (Jiang et al., 2022). One of the major challenges in kelp cultivation is dealing with the complex life cycle of these organisms that lack seeds. In the life cycle of kelps two free living, morphologically different multicellular bodies (generations) alternate; a haploid gametophyte and a diploid sporophyte. The sporophyte has a bladelike appearance and can reach meters in length. Haploid meiospores, released from reproductive sorus tissue on the adult sporophyte germinate and develop into filamentous, microscopic male or female gametophytes that colonize solid underwater surfaces. Spermatozoids released by the males are attracted to eggs exposed by the female gametophyte. After fertilization a sporophyte embryo develops, which initially is associated with the female gametophyte through the oogonial tube (Klochkova et al., 2019).

Sporophyte formation is the output of a sequence of multiple processes including vegetative gametophytic growth, gametogenesis, fertilization, embryogenesis, vegetative sporophytic growth, apical-basal polarity establishment and differentiation. A disturbance to any of these processes impacts (the abundance of) sporophyte formation. By adjusting the nutrient composition, light intensity and gametophyte density, the onset of gametogenesis, and thus the transition from a gametophyte into a sporophyte, can be induced in a laboratory setting (Aizo et al., 1984; Ebbing et al., 2025; Ebbing et al., 2021). Controlling and synchronizing early development, including the transition from gametophyte into sporophyte, is instrumental to the provision of high quality starting material for cultivation. As it turns out, microbiota also play a role in this process.

To identify the role of microbiota in kelp growth, Druehl & Hsiao, (1969) surface sterilized a piece of thallus that produces meiospores (the sorus) and allowed the released meiospores to germinate in the absence of a culturable microbiome. In these axenic incubations a brown mat developed on the bottom of the culture flasks, but no sporophytes formed. However, in uncleaned control incubations with an unknown microbiome (xenic cultures) vigorous sporophyte development was observed (Druehl & Hsiao, 1969). In addition to this work showing a putative obligatory symbiosis between kelp and associated microbiota, also facultative symbioses may affect the efficiency of kelp reproduction. Bacterial isolates added to xenic gametophytes either positively or negatively influenced sporophyte number (Park et al., 2023). It is evident that bacteria influence growth, development and reproduction of sugar kelp. However, it remains elusive which bacteria are the key players and on what developmental processes they take hold.

Using an adapted version of the method of Druehl et al. (1969), we were able to dissect the role of microbiota on *S. latissima*. We show that sugar kelp gametophyte cultures devoid of culturable bacteria display severely distorted early development and reproduction. In addition, the sporophytes that do form under axenic conditions are severely impaired. These defects can be partially or fully complemented by the addition of specific bacterial isolates from the sugar kelps core microbiome.

## Materials and methods

### Starting material and synchronization of meiospore release

Mature and reproducing *S. latissima* sporophytes on which developing sori were clearly visible by local darkening of the blade tissue were obtained from the Netherlands Institute for Oceanographic Research NIOZ; Texel, NL. From these sori square pieces of ∼ 2 × 2 cm were excised. The surface of excised fragments were cleaned thrice by rubbing with paper towels alternated by washing with filter sterilized artificial seawater (Instant Ocean) F/2 (FSF/2). From hereafter experiments were performed inside a flow cabinet using sterile tools and wearing nitrile gloves. The blade fragments were submerged in 1% (v/v) chlorine solution in FSF/2 for 1 minute after which they were washed 3 times 5 minutes in FSF/2. After the third wash, the outer 1 mm of the sorus fragments was removed using a sterile scalpel. Next, the sterilized fragment were dried by gently wiping with a tissue and left to dry in between sterile Whatman paper for approximately 14 hours at 10°C in the dark. Single sterilized and dried thallus pieces were transferred into 25 mL FSF/2 in a sterile 50 mL Falcon Tube and incubated on a rotary shaker at 10°C until meiospore release occurred. Meiospore release was evident when the previously clear and colorless medium became cloudy and brown after 5 to 15 minutes. After the water had turned brown and cloudy the thallus piece was removed using tweezers and the tube was placed on ice.

### Dilution of meiospore suspension

Elimination of putative remaining bacteria was performed by serial dilution of the sporulate. Tenfold dilution series until 10^9^ were made by pipetting 3 mL of the sporulate into 27 mL FSF/2. Meiospore density was determined in all dilutions using a hemocytometer. All experimentation was performed using the dilution step that contained 50-100 meiospores per incubation.

### Confirmation of microbial depletion

Of each individual meiospore incubation 50 µL of the medium was plated on solidified Sea Water Complete (SWC ATCC) medium, incubated at room temperature for 14 days and checked for absence of bacterial colonies. In addition, meiospores were allowed to germinate on F/2 medium solidified with 1% agar under standard vegetative gametophyte growth conditions (10°C, 14:10 L:D and 14 µmol/m^2^*s^-1^ red light). The resulting gametophyte cultures were microscopically examined regularly for the duration of one year. For all experiments describing axenic phenotypes, kelp cultures were used that did not yield colonies on both SWC and F/2 plates.

### Isolation of the culturable core microbiome

Bacteria were isolated from an established laboratory culture that had been growing in our laboratory for >5 years and from gametophyte cultures newly obtained from meiospores from Norway and the Netherlands, respectively. Sorus tissue from Norway was obtained from Hortimare BV, Heerhugowaard, NL. Cultures with reduced microbiome were obtained by using the last dilution serially diluted meiospore suspensions with normally developed gametophytes (see dilution of meiospore suspensions). Bacteria were isolated both from gametophytes growing in liquid F/2 and on solid F/2.

For isolation of all culturable bacteria, 5 media were used differing in carbon source as described by Lian et al. (2021). These media were prepared by adding 2 g/L casamino acids (CAS), 2 g/L succinate (SUC), 1 g/L yeast extract and 1 g/L peptone (YAP) or a *S. latissima* gametophyte extract based on 2 g/L wet weight to ESW (Enriched Sea Water) 2 g/L glucose (GLU). Gametophyte extract was made by crushing washed gametophytes using glass beads and filter sterilizing the suspension twice by pressing it through 0.2 um filters. After isolation on specific media, all bacterial strains were transferred onto standard SWC medium on which all of them grew sufficiently well for further experimentation.

### Isolation of the culturable core microbiome

Bacteria were isolated from established *S. latissima* gametophyte cultures that had been growing in our laboratory for >5 years and from gametophyte cultures newly obtained from meiospores from Norway and the Netherlands, respectively. Sorus tissue from Norway was obtained from Hortimare BV, Heerhugowaard, NL. Cultures with reduced microbiome were obtained by using the last dilution serially diluted meiospore suspensions with normally developed gametophytes (see dilution of meiospore suspensions). Bacteria were isolated both from gametophytes growing in liquid F/2 and on solid F/2.

For isolation of all culturable bacteria, 5 media were used differing in carbon source as described by Lian et al. (2021). These media were prepared by adding 2 g/L casamino acids (CAS), 2 g/L succinate (SUC), 1 g/L yeast extract and 1 g/L peptone (YAP) or a *S. latissima* gametophyte extract based on 2 g/L wet weight to ESW (Enriched Sea Water) 2 g/L glucose (GLU). Gametophyte extract was made by crushing washed gametophytes using glass beads and filter sterilizing the suspension twice by pressing it through 0.2 um filters. After isolation on specific media, all bacterial strains were transferred onto standard SWC medium on which all of them grew sufficiently well for further experimentation.

### Bacterial characterization by 16S gene sequencing

Bacterial colonies picked from the agar plates were directly used as a template in PCR amplification of the 16S rRNA gene using the 27f forward primer and the 1390r reverse primer (Mao et al., 2012). Amplified fragments were sequenced (Macrogen, Netherlands using Sanger sequencing) and manually curated sequences (using SnapGene Viewer) were blasted against type material using MegaBLAST. Based on the 98.65% sequence similarity threshold for 16S sequences (Kim et al., 2014) we determined all but one isolate to species level.

### Screening the effects of specific bacterial isolates on growth and development of *S. latissima*

Axenic meiospores were brought into FSF/2 medium with a predicted density of 60 meiospores in 2 mL culture medium in 24 wells plates (Sarstedt) and left to settle for 3 days before introducing bacteria. Bacterial suspension cultures were grown in liquid SWC until an OD_600_ of approximately 1, centrifuged at 2000xg for 5 minutes and resuspended in FSF/2 after which they were diluted and added to the axenic meiospores. Based on OD_600_ measurements with the assumption that all bacteria had a size similar to *E. coli* (where OD_600_ of 1 corresponds to ∼ 8*10^8^ cells/ml), a calculated 800 bacterial cells per mL were added to settled axenic meiospores by replacing 100 µL of the culture medium with 100 µL of bacterial suspension. For the negative controls 100 µL of the culture medium was replaced by 100 µL fresh FSASWF/2. For each inoculation 3 replicates were used and for the negative control 32 replicates.

### Microscope settings and image analysis

Image acquisition for the microbiome complementation experiments was performed on a Stellaris DMI-8 confocal microscope (Leica Microsystems) controlled with LAS X 4.7.0.28176 software, using a 10x 0.4 NA HC PL APO objective and 600Hx scan speed. Samples were excited with 492nm laser light using a white light laser (2%), and autofluorescence images were collected using a 681-750nm emission filter on a HyD S detector with pinhole dimensions set to 4 AU. Black and white images were collected using the trans PMT channel. 5×5 (x*y) images were collected per well of a 96 wells plate and subsequently combined to a single image using the stitching function of the LAS X software. Imaging of *Saccharina latissima* colonies and individual sporophytes was performed using a Nikon Diaphot 200 inverted microscope with a Ds Fi1 Camera (Nikon) and 10x Nikon MC1 0.25 NA LWD, 20x Nikon MC2 0.4 NA or 40x Nikon MC3 0.55 NA objectives.

Growth of gametophytes was scored after 30 days by acquisition of autofluorescence and trans PMT images of all treatments. Number of individuals, the area and the solidity were determined using ImageJ version 1.51f (http://imagej.nih.gov/ij) (fig. S1) (as described by Vidali et al. (2010).

All overlapping and partially visible colonies at the border of an assembled 5×5 fields of view image were excluded from the dataset. To study the differential effect on the 2 sexes, male and female colonies were separated based on filament size and colony morphology where male filaments are much smaller compared to females (Bi & Zhou, 2014).

### Statistical methods

Statistical analyses were performed in R version 4.5.1. Shapiro-Wilkinson tests were used to test for normality. On the data of the area and solidity of gametophytes we performed a Shapiro wilks to test for normality. After determining that all the data was not normally distributed we performed a Kruskal Wallis test whereafter we performed a post hoc Dunn test with holm multiple testing correction. To analyse the data on bacteria induced oogenesis and meiospore germination we conducted a pairwise chi-square test with holm statistical adjustment. For data that were not normally distributed, we used Kruskal Wallis tests to determine significance whereafter we performed a post hoc Dunn test with Holm multiple testing correction. A statistical significance of 0.05 was chosen for all tests.

## Results

### Axenic gametophytes of *Saccharina latissima* are viable

To obtain axenic gametophyte cultures we released meiospores from surface sterilized sorus and additionally diluted the sporulate to eliminate all microbiota. Because no bacterial growth was observed when the sporulate was cultured on rich medium and no bacteria were observed while monitoring gametophyte development we deemed these cultures as axenic. The fraction of meiospores that germinated under axenic conditions did not differ from the xenic control which constituted of uncleaned meiospores still containing a microbiome from the parental sporophyte (fig. S1). Meiospore germination thus not relied on the presence of bacteria. However, non-germinated meiospores persisted in axenic cultures compared to xenic cultures where (remnants of) non-germinated spores did decay (fig. S1).

### Axenic *S. latissima* gametophytes show reduced vegetative growth, spontaneous oogenesis and distorted development of sporophytes

To test whether the absence of a microbiome affected growth and development of *Saccharina latissima*, gametophytes were cultured on solid marine agar medium, which allowed us to track individual gametophytes. Additionally, potential remaining bacteria were immobilized on solid medium and bacterial growth in xenic cultures could be observed as colonies that formed on the plate (blue arrows figure 1). After 22 days, clear and consistent phenotypic differences between axenic and xenic cultures were evident (figure 1). In xenic cultures, both male and female meiospores had developed into gametophytes consisting of elongated, branching cell files (Figure 1a and b). Long term (8 months) growth of xenic gametophytes on solid marine agar medium showed clear, dark-brown pigmented colonies (figure 1c and d) with clear filaments of cells branching outwards. These phenotypes constitute healthily growing gametophytes.

**Figure 1.**
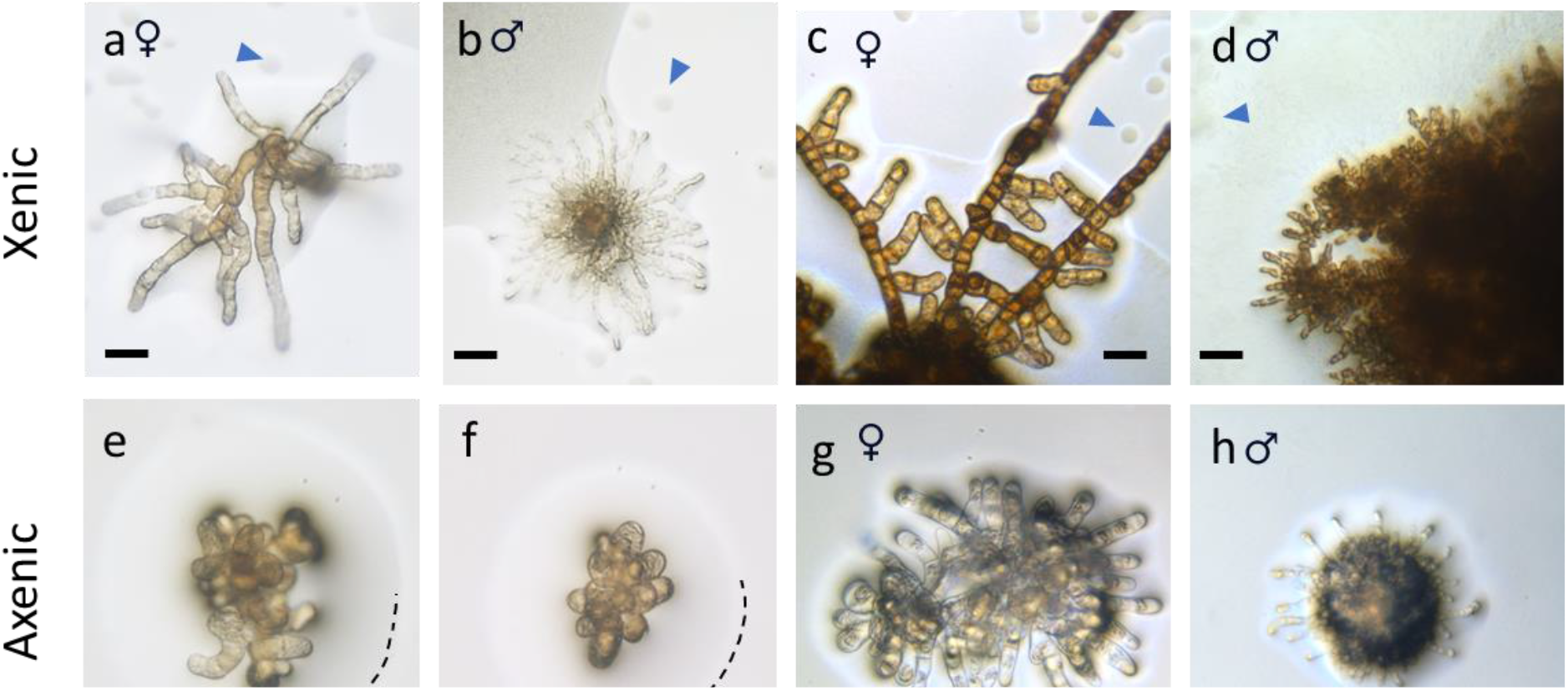
*S. latissima* gametophytes growing vegetatively on solid marine agar in the presence and in the absence of visible bacterial growth. Images a-d show xenic gametophytes from uncleaned meiospores and e-h show gametophytes from axenic meiospores. Images a, b, e and f were taken 22 days after germination and images c, d, g and h were taken after ∼ 8 months. Note that xenic gametophyte colonies grew outwards with elongated cells and remained growing and developing for at least 8 months (a-d, blue arrowheads point towards bacterial growth). The dashed line in e and f shows part of the water film that adheres onto the gametophytes, but no bacterial colonies are visible. Axenic gametophyte colonies failed to grow outward, grew slower compared to the xenic cultures and had died after 8 months (g, h). The scale bar represents 25 µm. Images are representative images of 255 and 57 colonies for xenic and axenic colonies, respectively.

In contrast, axenic gametophytes were smaller, and lacked filamentous outgrowths (Figure 1e and f). After 8 months, cells of axenic gametophytes had disintegrated and the remains were found collapsed inside the persisting cell wall, indicating that these colonies were dead (Figure 1g and h). Interestingly, the dead gametophytes had not decayed, indicating that bacteria that degrade *Saccharina latissima* were not present. Thus, the absence of a culturable microbiome severely impacted gametophyte growth and morphogenesis.

Next, we studied gametogenesis and sporophyte development in both xenic and axenic cultures. Because these processes were not observed on marine agar, possibly because egg protrusion and fertilization depend on submergence in water, we studied these processes in liquid medium. We decided to focus on gametogenesis in females and not in males mainly due to larger cell size and the sessile nature of eggs as opposed to motile sperm, which made females more practical to study. We found that oogenesis and sporophyte development is severely impacted in axenic gametophytes when compared to xenic cultures (figure 2). Even when cultivated under conditions that typically suppress gametogenesis and keep gametophytes growing vegetatively (red light without the addition of iron to the cultivation medium, (see methods)), female axenic gametophytes showed abundant spontaneous gametogenesis (figure 2e, eggs/zygotes orange arrowheads and f, empty oogonia blue arrowheads) unlike the xenic cultures. Cells of axenic gametophytes were unusually elongated when compared to those in xenic cultures, as are the oogonia that remain after egg protrusion (figure 2f, g, c, blue arrowhead).

**Figure 2.**
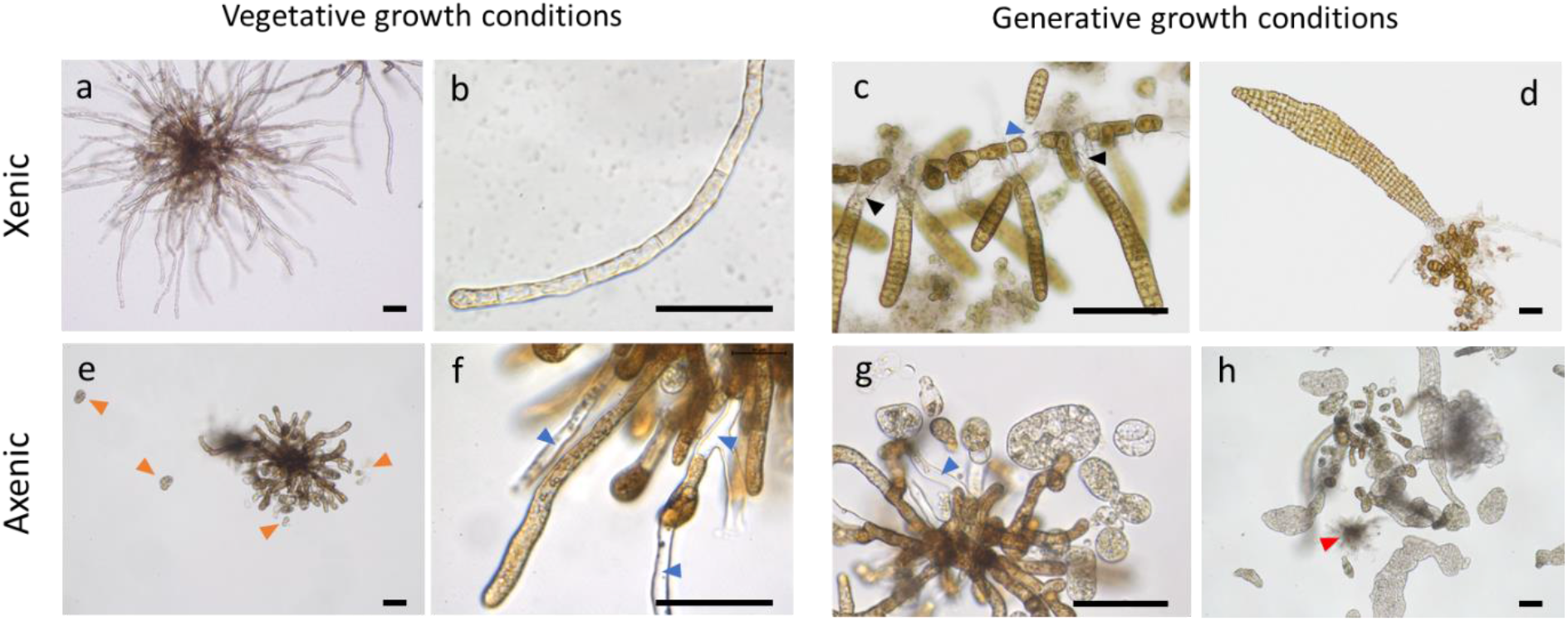
Gametogenesis and sporophyte development of *S*. in the presence and in the absence of a microbiome. The top row (a-d) shows xenic gametophytes with an undefined microbiome from the parent plant cultivated under conditions that promote vegetative (a and b) and generative (c (1 week) and d (3 weeks)) growth. The bottom row shows axenic gametophytes also cultured under vegetative (e and f) and generative (g (1 week) and h (3 weeks)) conditions. Note spontaneous gametogenesis in axenic gametophytes even under vegetative conditions (eggs shown by orange arrowheads in e and empty oogonia by blue arrowheads in f). red and black arrowheads point towards male gametophyte and rhizoids of sporophytes, respectively. The scale bar represents 25 µm.

One week old axenic sporophytes were detached from the oogonium, lacked polarity and rhizoids, and their poorly defined cells showed abnormally weak pigmentation (figure 2 g compared to c). Xenic sporophytes of the same age remained attached to the oogonium, show the onset of rhizoid formation, had well defined cells and proper dark pigmentation (figure 2c). 3 week old axenic sporophytes continued to grow but the vast majority failed to establish polarity and the growth defects observed earlier were not overcome (figure 2h compared to d).

### Isolation and characterization of the microbiome of *S. latissima* gametophytes

Using a diversity of bacterial culture media, 10 bacteria associated with kelp cultures were isolated and subsequently characterized based on colony characteristics and the 16S rRNA gene sequence. For 9 out of 10 isolates we found a hit in the NCBI database with the percentage identity exceeding the species identification threshold of 98.65% (Kim et al., 2014) and we named them *C. baekdonensis, W. faeni, S. donghicola, V. titanicae, S. todarodis, A. stellipolaris, M. alkaliphilus, P. flavimaris* and *S. stellata* (table 1). For our *Erythrobacter* sp. isolate we stuck to naming it as a species within the genus *Erythrobacter* because the highest percentage identity of 98.53% remained under the threshold.

**Table 1.**
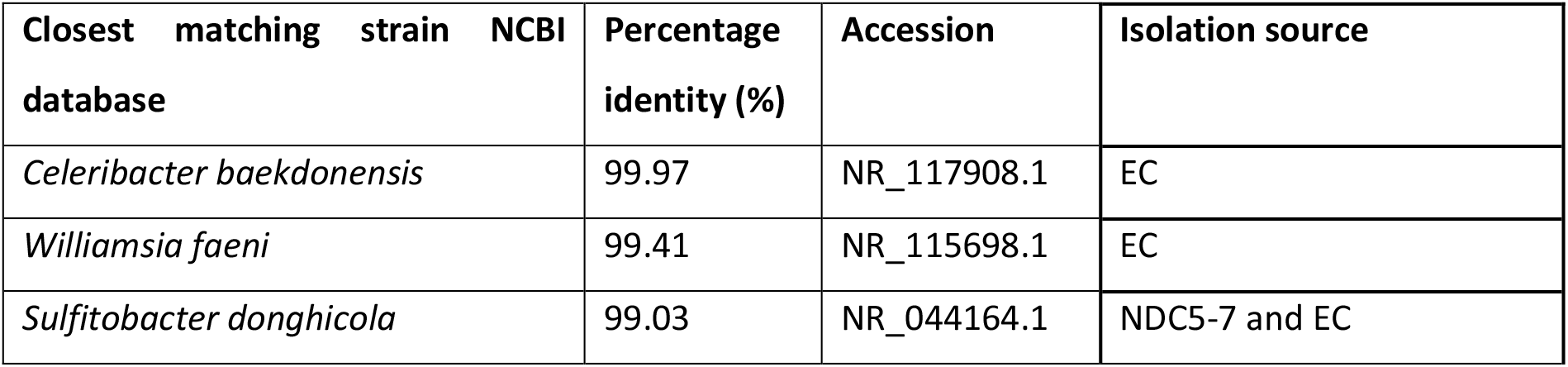
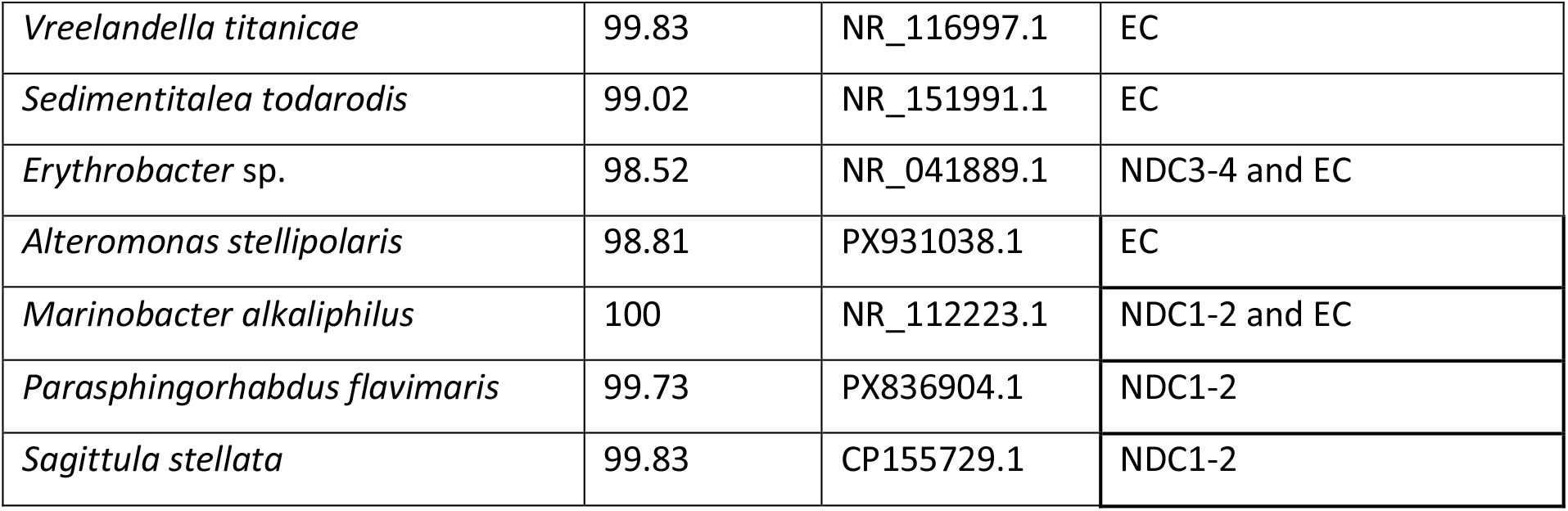
Identification of 10 bacterial isolates obtained from various laboratory cultures of *Saccharina latissima*. Bacteria were isolated from both established (EC) and new diluted (NDC) cultures that both showed healthy gametophyte growth.

| Closest matching strain NCBI database | Percentage identity (%) | Accession | Isolation source |
| --- | --- | --- | --- |
| *Celeribacter baekdonensis* | 99.97 | NR_117908.1 | EC |
| *Williamsia faeni* | 99.41 | NR_115698.1 | EC |
| *Sulfitobacter donghicola* | 99.03 | NR_044164.1 | NDC5-7 and EC |
| *Vreelandella titanicae* | 99.83 | NR_116997.1 | EC |
| *Sedimentitalea todarodis* | 99.02 | NR_151991.1 | EC |
| *Erythrobacter* sp. | 98.52 | NR_041889.1 | NDC3-4 and EC |
| *Alteromonas stellipolaris* | 98.81 | PX931038.1 | EC |
| *Marinobacter alkaliphilus* | 100 | NR_112223.1 | NDC1-2 and EC |
| *Parasphingorhabdus flavimaris* | 99.73 | PX836904.1 | NDC1-2 |
| *Sagittula stellata* | 99.83 | CP155729.1 | NDC1-2 |

We isolated bacteria from two sets of healthy growing gametophyte cultures. From one established culture (EC) that had been maintained in the laboratory for more than 5 years and from 7 new cultures grown from serially diluted meiospores (NDC1-7). NDC1-2 and NDC3-7 originated from different parent plants from the same location (Texel the Netherlands).

Most diverse isolates were obtained from the established and undiluted *S. lattissima* culture (8 species). *P. flavimaris* and *S. stellata* were isolated exclusively from new diluted cultures (NDC1-2) while *C. baekdonensis, W. faeni, V. titanicae, S. todarodis* and *A. stellipolaris* were only found in the established culture (EC) (table 1). *M. alkaliphilus, Erythrobacter* sp. and *S. donghicola* were found in both established and new, diluted cultures. New, diluted cultures had a lower microbial diversity compared to the established culture. From new, diluted cultures we isolated either only *Erythrobacter* sp. (NDC3-4) or *S. donghicola* (NDC5-7) or the combination of *M. alkaliphilus*., *P. flavimaris* and *S. stellata* (NDC1-2).

### Growth of male and female gametophytes is restored after complementation with a wide variety of bacteria, with a different response in males and females

To test the effect of the different isolated bacteria on the growth and development of *Saccharina latissima* gametophytes, we germinated axenic spores and reintroduced specific bacterial isolates. Besides the bacteria isolated from *S. latissima*, we introduced two bacterial strains previously shown to promote healthy growth and development in the green macroalga *Ulva compressa*; *Roseovarius* sp. strain MS2 and *Maribacter* sp. strain MS6 (Spoerner et al., 2012). Since male and female gametophytes of *Saccharina latissima* have a different colony morphology, we were able to separate the effects of bacteria on both groups. To compare the effect of bacteria on gametophyte size and morphology we measured gametophyte area and solidity. Gametophytes with a relatively low solidity are better able to form outgrowing filaments compared to gametophytes with a relative high solidity that have a clumpy morphology (figure S2 for quantification visualization). These shape descriptors discriminate between the typical shapes associated with axenic and xenic gametophyte cultures (figure 1).

In females the addition of *W. faeni, S. donghicola, S. stellata, Roseobacter* sp., *M. alkaliphilus*., *V. titanicae, Erythrobacter* sp., *C. baekdonensis* and *Roseovarius* sp. led to an increased area and a reduced solidity, correlating with bigger gametophytes with more filamentous growth when compared to the axenic control (figure 3A, B). Addition of the other 3 tested isolates *P. flavimaris, A. stellipolaris* and *Maribacter* sp. MS6 did not significantly affect size and filamentous outgrowth in females when compared to the axenic control.

**Figure 3.**
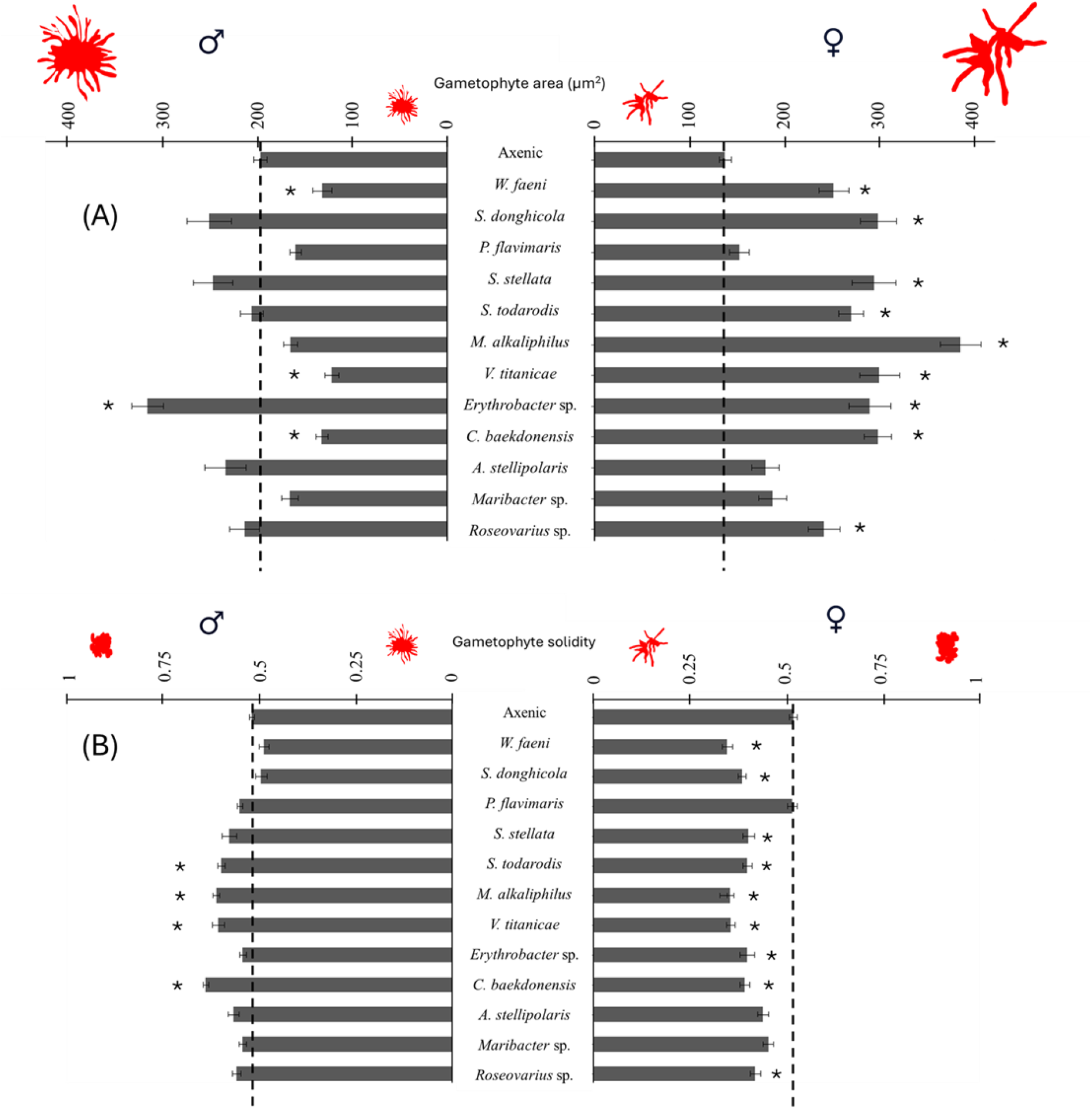
Gametophyte colony size and shape are affected by the addition of specific bacterial isolates when compared to an axenic control. Colony size is determined by measuring the surface area of the colonies in square micrometers (A). The shape descriptor solidity is calculated by dividing the area in A over the area of the convex hull area and is a measure for outgrowth of colonies (B) (figure S2 for methods visual). The dotted line in all graphs shows the control, which constitutes of axenic gametophytes and aids in visual comparison. Asterisks indicate significant differences compared to the axenic control (p=0.05, N=3, n=90).

**Figure 4.**
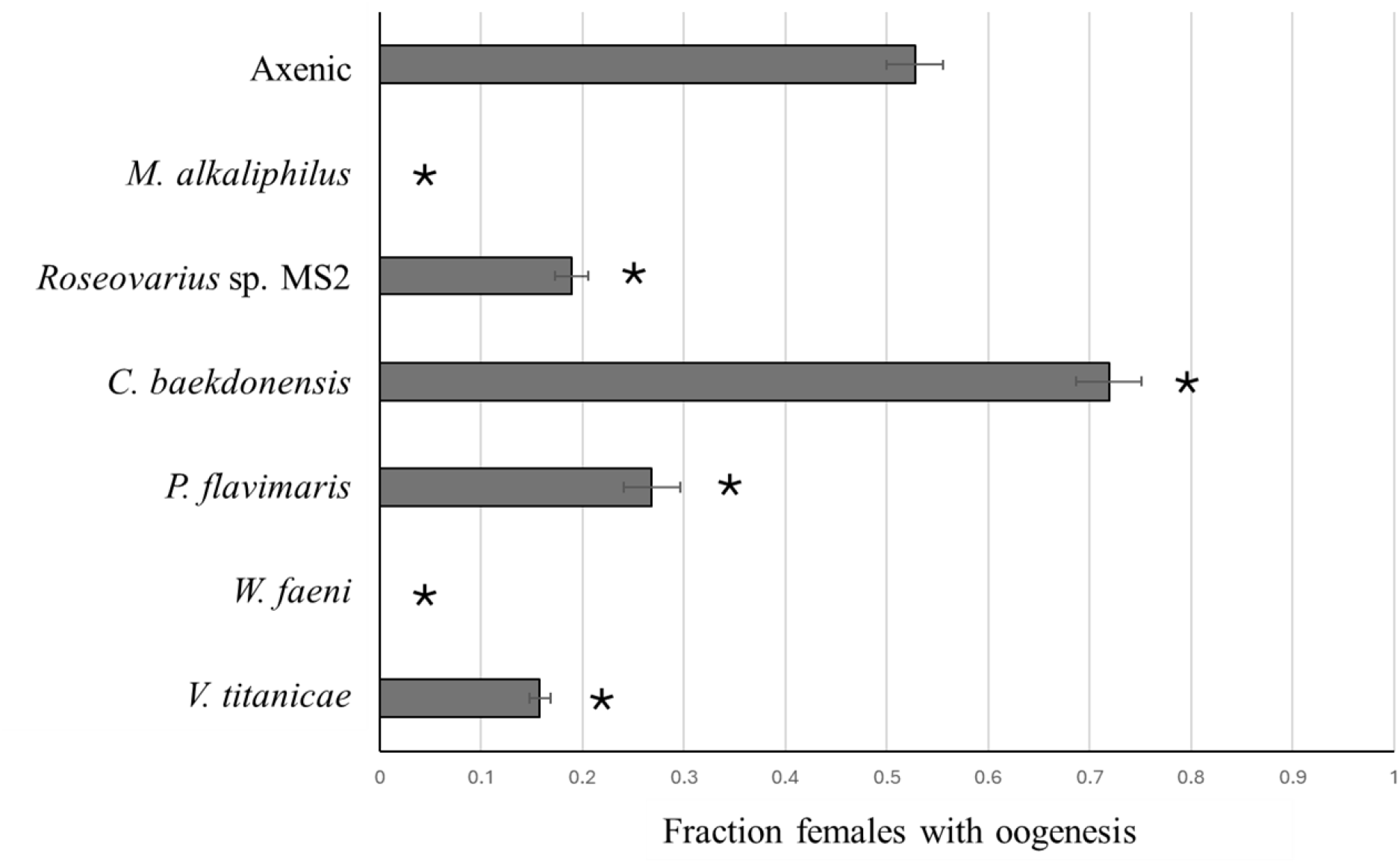
Bacterial isolates stimulate or suppress oogenesis in female gametophytes when compared to axenic control. Colonies with eggs, young sporophytes and/or empty oogonia were counted and divided over the total number of colonies, showing the fraction of females that had undergone oogenesis under vegetative growth conditions (N=4, n=35-53). Asterisks show significant differences with axenic incubations where α is equal to or lower than 0.05.

The same bacterial isolates had a different effect on the size and shape of male gametophytes. The addition of *V. titanicae* and *C. baekdonensis* to males negatively influenced size and solidity, resulting in smaller and more clumpy gametophytes when compared to the axenic control (figure 3A, B left). In males the addition of *Erythrobacter* sp. and *W. faeni* respectively positively and negatively influenced growth, but without significantly affecting the solidity. *S. todarodis* and *M. alkaliphilus* reduce solidity but not size leading, disturbing filamentous outgrowth while not affecting growth.

Not all bacterial isolates affected size and or filamentous growth in gametophytes. The addition of *P. flavimaris, A. stellipolaris* and *Maribacter* sp. MS6 had no effect on size and solidity in neither male nor female when compared to their respective controls (figure 3A, B).

### Different bacterial isolates have either positive or negative effects on oogenesis

Here we set out to quantify oogenesis under axenic conditions and in the presence of bacterial isolates. We found that under axenic conditions about half of the female gametophytes underwent oogenesis (figure 3). Addition of *M. alkaliphilus* and *W. faeni* completely suppressed oogenesis and *Roseovarius* sp. MS2, *P. flavimaris* and *V. titanicae* significantly reduced oogenesis when compared to axenic incubations (p<0.004, p<0.004, p<0.004, p=0.04 and p<0.004, respectively). On the contrary, the addition of *C. baekdonensis* stimulated oogenesis when compared to the axenic incubations (p=0.05).

### Abnormal shape, pigmentation and size observed in axenic sporophytes was restored by complementation with the majority of the bacterial isolates from *Saccharina latissima*

Sporophyte development under axenic conditions and in coculture with bacterial isolates was qualitatively assessed in the mixed sex cultures of the previous experiment. Consistent with earlier observations (figure 2g, h), we observed that under axenic conditions sporophytes did form in all incubations, but that they generally failed to develop a clearly defined apical-basal axis, lack dark pigmentation and did not grow longer than 500 µm in the 4 week growth period (table 2, figure 5a). Additionally, eggs were often extruded as beads on a string instead of a single spherical cell and released into the medium rather than remaining attached to the oogonial tip (figure 5a and S3).

**Table 2.** Qualitative comparison of sporophyte development under axenic conditions and in coculture with single bacterial isolates. Rows indicate whether or not sporophytes had formed, sporophyte size, pigmentation and filamentous outgrowths of thallus margins. + indicates that this was observed in all incubations (20 replicates for axenic incubations and 4 replicates per cocultivation), – indicates it was found in none of the incubations and a +/-indicates it was found in some, but not all incubations. Green and red indicate normal and abnormal development, respectively for all observations and orange indicates a variation of normal and abnormal development in the replicates of that treatment. Qualitive assessment is extracted from the images shown in figures S3 and S4.

|  | Axenic | *Williamsia* sp. | *Sulfitobacter* sp. | *Sphingopyxis* sp. | *Sagittula* sp. | *Roseobacter* sp. | *Marinobacter* sp. | *Halomonas* sp. | *Erythrobacter* sp. | *Celeribacter* sp. | *Alteromonas* sp. | *Roseovarius* sp. | *Maribacter* sp. |
| --- | --- | --- | --- | --- | --- | --- | --- | --- | --- | --- | --- | --- | --- |
| Sporophytes formed | + | +/- | + | + | + | + | +/- | + | + | + | + | + | + |
| Sporophytes > 500 µm | - | + | + | +/- | + | + | + | + | + | + | + | + | - |
| Sporophytes with dark pigmentation | +/- | + | + | +/- | + | + | + | + | + | + | + | + | +/- |
| Sporophytes with filamentous outgrowths | - | - | - | - | - | +/- | - | - | - | - | - | - | - |

**Figure 5.**
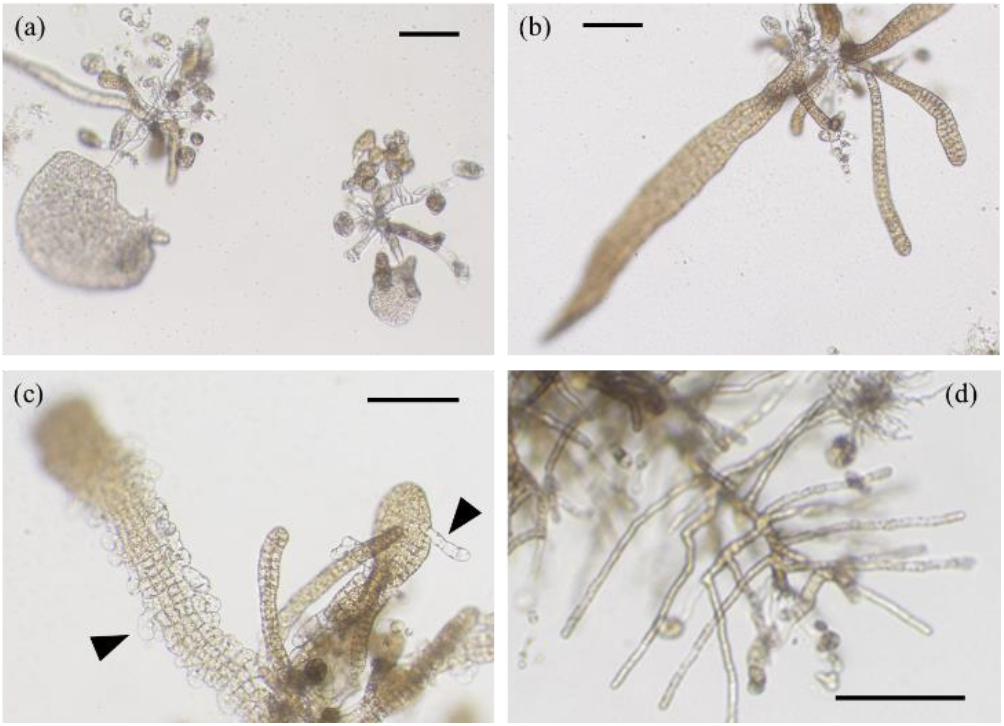
Diverse sporophytes form when developing axenic or in the presence of specific bacterial isolates. Axenic sporophytes (a) with relatively small, pale, isodiametric thalli. Sporophytes in cocultivation with *Erythrobacter* sp. (b) with dark, long and relatively large thalli when compared to axenic sporophytes, representative for normal development. Sporophytes developed filamentous outgrowths from the thallus margins in some incubations with *Sedimentitallea todarodis* (black arrowheads in c) and *Williamsia faeni* and *M. alkaliphilus* prevented sporophyte development in some but not all of the incubations (d image of cocultivation with *M. alkaliphilus*). Scale bar represents 100 µm. For whole panel of images see figure S4.

When coculturing *Saccharina latissima* gametophytes with either *S. donghicola, P. flavimaris, S. stellata, V. titanicae, Erythrobacter* sp., *C. baekdonensis, A. stellipolaris* or *S. todarodis* elongated, dark pigmented sporophytes bigger than 500 µm developed after 4 weeks (table 2, figure 5b). These sporophyte characteristics are generally observed in xenic cultures and constitute normal development (figure 2 d).

Addition of *Maribacter* sp. and *P. flavimaris* yielded sporophytes with similar characteristics as axenic sporophytes. Sporophytes were consistently small in all replicates of *Maribacter* sp., but in half of incubations with *P. flavimaris* sporophytes bigger than 500 µm were observed. The addition of *M. alkaliphilus* and *W*. sp. suppressed sporophyte formation in some of the incubations, but when sporophytes did form in the presence of these bacteria they did grow bigger than 500 µm and developed dark pigmentation (figure 5d). When cocultured with *S. todarodis* filamentous outgrowths emerged from peripheral cells of the sporophyte thallus in half of the incubations (figure 5c).

## Discussion

Intimate symbiosis between seaweeds and associated microbiota have amongst others been demonstrated in model species for green and brown seaweeds *Ulva compressa* (Chlorophyceae) and *Ectocarpus siliculosus* (Phaeophyceae) (Spoerner et al., 2012; Tapia et al., 2016). In these macroalgae with a relatively simple body plan, consisting of a flat or tubular thallus with basal rhizoids (*U. compressa*) and branching cell files (*E. siliculosus*), a wide variation of bacteria from several different families have been isolated that orchestrate developmental processes including cell division, cell elongation and rhizoid formation. Here we isolated bacteria from the sugar kelp (*Saccharina latissima*, Phaeophyceae) microbiome of which some isolates similarly affect cell division and filamentous outgrowth, but additionally influenced oogenesis and pigmentation in a seaweed with a complex life cycle, organogenesis and embryogenesis.

Studying the effects of the complex microbiome on growth and development by inoculation of isolates to axenic cultures is essentially creating a minimal system set-up. We were able to isolate and test a total of 10 bacterial strains from our kelp cultures. This set has a rather low diversity when compared to for instance the 47 bacteria used by Grueneberg et al. (2016) to complement axenic *U. compressa* cultures, but similar to the 9 isolates used by Tapia et al. (2016) to complement axenic *E. siliculosus*. Interestingly, Grueneberg et al. isolated bacteria from coastal water samples, whereas Tapia et al. and we isolated the bacteria from laboratory cultures. Despite the lower microbial diversity isolated from laboratory cultures, the relative abundance of morphogenetic bacteria was much higher: 8/10 for this study and 6/9 for Tapia et al., compared to 14/47 in the study of Grueneberg et al. (2016). These observations are in line with the core microbiome hypothesis (Neu et al., 2021), where microbial key players are likely to be associated tightly with the seaweed and likely to persist after dilution, increasing the relative abundance of bacteria that would affect growth and development in cultures with a reduced microbiome. When mining for morphogenetic bacteria, healthy growing laboratory cultures with a reduced microbiome appear to be an efficient source.

One genus of morphogenetic bacteria that we isolated from *Saccharina latissima* overlaps with genera found in *Ectocarpus* sp.. In *Ectocarpus* sp. *Marinobacter* sp. has filament inducing activity, leading to longer filaments growing outward from the colony. Axenic *Ectocarpus* sp. colonies grow into clumpy structures (Tapia et al., 2016). We found that *Marinobacter alkaliphilus* has a similar positive effect on female gametophytes of *Saccharina latissima*, but negatively influences male development. Female gametophytes are larger and show more filamentous outgrowth than the axenic controls while male gametophytes unaffected in size compared to axenic gametophytes, and show less polar outgrowth from the center of the colony (figure 3).

We found an additional 7 isolates with growth promoting effects on female gametophytes (*Celeribacter baekdonensis, Sagitulla stellata*., *Sulfitobacter donghicola, Sedimentitalea todarodis* (all Rhodobacteraceae), *Williamsia faeni* (Williamsiaceae), *Erythrobacter* sp. (Erythrobacteraceae), and *Vreelandella titanicae* (Halomonadaceae)). Female gametophytes incubated with bacteria from these genera showed similar phenotypes as the ones with growth promoting effects in *Ectocarpus* sp.. Similar to *M. alkaliphilus* and *S. todarodis*, three of our isolates (*Sagitulla stellata, S. donghicola* and *W. faeni*), male gametophytes were unaffected in growth. Like *S. todarodis, C. baekdonensis* had a negative effect on male gametophyte growth, resulting in smaller gametophytes. *Erythrobacter* sp. has a growth promoting effect on both male and female cultures, indicating that the processes *Erythrobacter* sp. influences were not male-female specific, but of a more general nature. *Erythrobacter* sp. is therefore a good candidate to setup with gametophytes in a coculture with a minimal but sufficient microbiome.

Besides the growth promoting bacteria, two additional bacterial genera were consistently isolated that had no effect on either male or female gametophyte growth (*Alteromonas stellipolaris* and *Parasphingorhabdus flavimaris*). Their role is unknown, however, *A. stellipolaris* broadly associates with brown seaweeds as it is a known pathogen of *Saccharina japonica* (Peng & Li, 2013) and was also isolated from the *Ectocarpus* sp. microbiome, where it also did not influence growth (Tapia et al., 2016).

Spontaneous gametogenesis observed in axenic females led us to hypothesize that the process of oogenesis could be regulated by microbiota. Under non-inducing conditions, we quantified egg production in axenic and cocultured gametophytes and found that in axenic cultures half of the females produced eggs. *W. faeni* and *M. alkaliphilus* fully suppressed oogenesis, while these isolates also had the strongest positive effect on vegetative growth of females suggesting that these bacteria play a role in the transition between vegetative and generative growth in sugar kelp. One of the bacterial strains tested here, *C. baekdonensis*, has a promoting effect on gametogenesis in females. It would be very interesting to obtain more detailed mechanistic insight in this effect, since it could provide a way to synchronize egg production and thus sporophyte formation for mariculture. Five bacterial strains show a negative effect on female gametogenesis, which suggests the intriguing possibility of inhibiting gametogenesis. Tentatively, this could be employed in a system where gametophytes are efficiently bulked up in the presence of a gametogenesis-inhibiting microbiome, after which the microbiome can be exchanged for a gametogenesis-promoting microbiome to stimulate the formation of the economically important sporophyte.

In contrast to *Ectocarpus* sp., the sporophyte of *Saccharina latissima* is a large and complex plant with tissue differentiation and various distinct organs (Boscq et al., 2025; Theodorou & Charrier, 2023). Studying the role of bacteria in growth and development of *Saccharina latissima* is therefore not restricted to monitoring colony shapes, but also involves their role in the development of the complex sporophyte. Embryogenesis in Laminariales takes place on top of the female gametophyte and pelagic bacteria have access to the developing embryo. This, in combination with the severe phenotypes of axenic sporophytes that we observed suggests that bacteria play a role in the establishment of apical-basal polarity, regulating the direction of cell division, differentiation and polystromatisation (Peng & Li, 2013).

In axenic cultures sporophyte development is limited to amorphous, bleached sheets that lack normal cell patterning and have fewer rhizoids. This suggests that, among others, an apical-basal polarity axis fails to form. The lack of an apical-basal axis and fewer rhizoids is also observed in partheno-sporophytes of related kelp species *Undaria pinnatifida* and in sporophytes that have detached from their oogonium in early development (Dries et al., 2024). We established axenic cultures from the meiospore stage and because gametophytes show growth defects, the identification of the primary cause of the observed defects in sporophyte development is obscured. We did find however, that that all of the studied bacterial isolates (table 1) at least in part restored apical-basal patterning, pigmentation and rhizoid formation.

Though our results point to a seemingly very redundant and opportunistic symbiosis between diverse microbiota and *Saccharina latissima*, not all bacteria can restore gametophyte and sporophyte development; the *Ulva compressa* isolate *Maribacter* sp. MS6 (Spoerner et al., 2012) fails to do so.

In conclusion, we have found that a core microbiome is necessary for *Saccharina latissima* vitality, growth and development. The effect of the isolated bacteria differs on male and female gametophytes. Sporophyte development also depends on the presence of bacteria, and sporophyte development can, at least partially, be rescued by the same bacteria as promote gametophyte growth. The exact processes *Saccharina latissima* microbiome bacteria influence and the signaling molecules they use are a topic of future research. Elucidating these will provide a step towards developing a probiotic application for commercial *Saccharina latissima* cultivation.

## Acknowledgements

We would like to thank Gabriel Montecinos Arismendi from Hortimare BV and Klaas Timmermans and Robert Twijnstra from the national oceanographic institute and center of expertise in the Netherlands for the ocean, sea and coast (NIOZ, NL) for the provision of starting material for experiments. We thank Thomas Whichard from the University of Jena (DE) for the *Ulva compressa* bacterial isolates MS2 and MS6 and Chris Roelofsen (CDB) for technical assistance. Finally we would like to thank Wageningen light microscopy center (WLMC) for the use of their equipment and microscope facilities.

## Author contributions

OPL contributed to the investigation, methodology, visualisation, writing – original draft, and writing – review and editing. PACG contributed to the investigation, methodology, and writing – review and editing. DSe contributed to data analysis and visualisation. TK, DSi and OPL: conceptualisation and writing – review and editing.

## Supplemental figures

**Figure S1.**
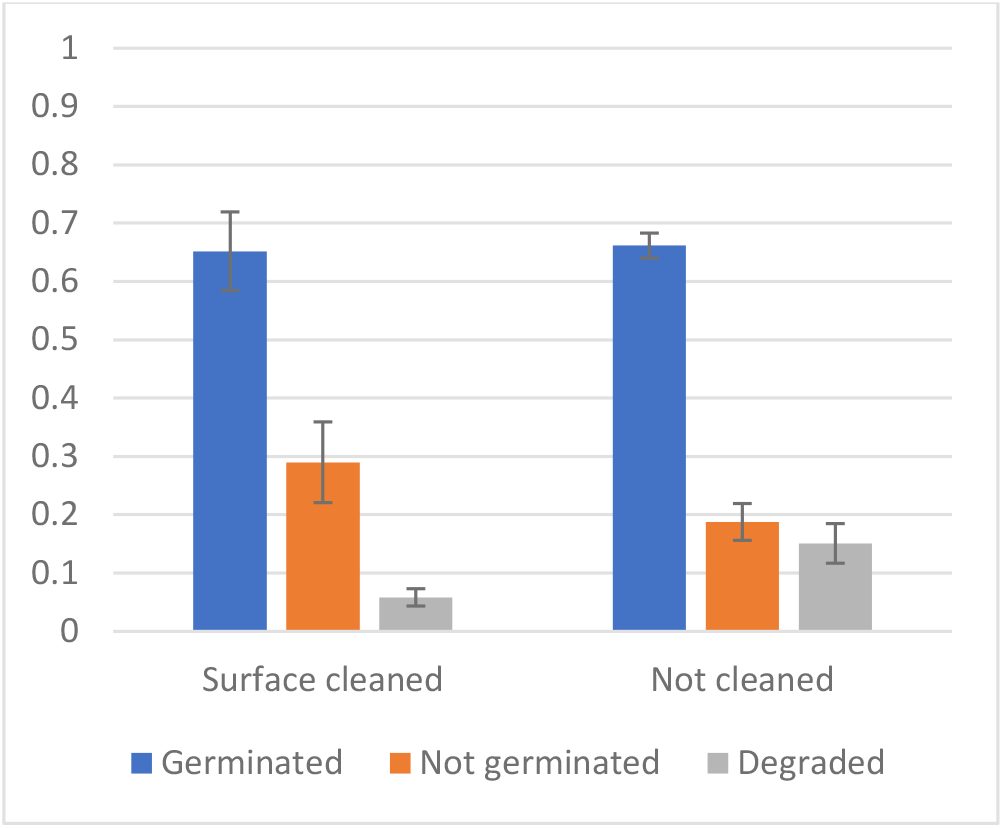
Meiospore germination is unaffected by surface sterilization methods. Surface sterilization does not significantly reduce meiospore germination when compared to uncleaned control sorus after 6 days (p=0.5786). The fraction of degraded meiospores, however, is lower and consequently the fraction of intact but not germinated meiospores is higher after surface cleaning when compared to untreated sorus (p=0.0053 and p=0.0003, respectively).

**Fig S2.**
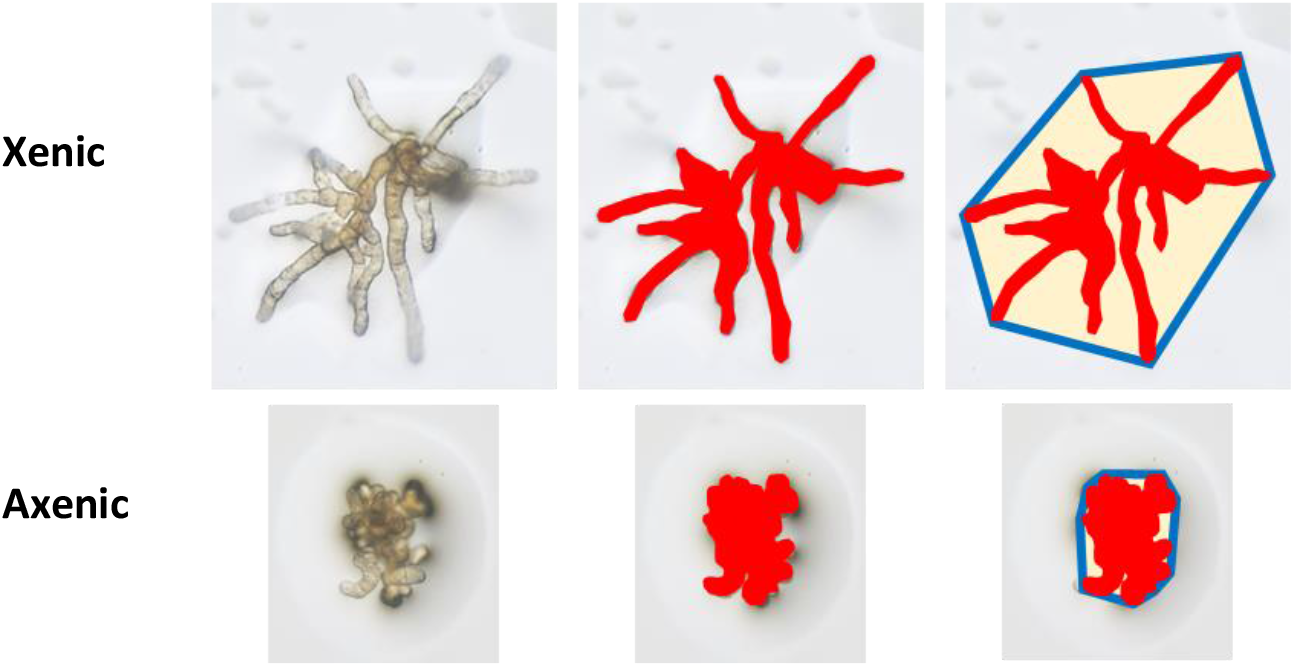
Area and solidity as variables to quantitatively compare growth and development of *S. latissima* gametophytes cultured with or without specific bacterial isolates. Of this female gametophyte both area (red) and convex hull (blue) were determined. The area is never larger than the convex hull area of the gametophyte, thus dividing the former over the latter yields a number smaller than 1; the solidity of the gametophyte. The solidity is a descriptor of shape where gametophytes with extended, outgrowing filaments (solidity < 0.5) are easily distinguished from gametophytes with clumpy growth (solidity > 0.5). Sizes of gametophytes are relative to each other.

**Figure S3.**
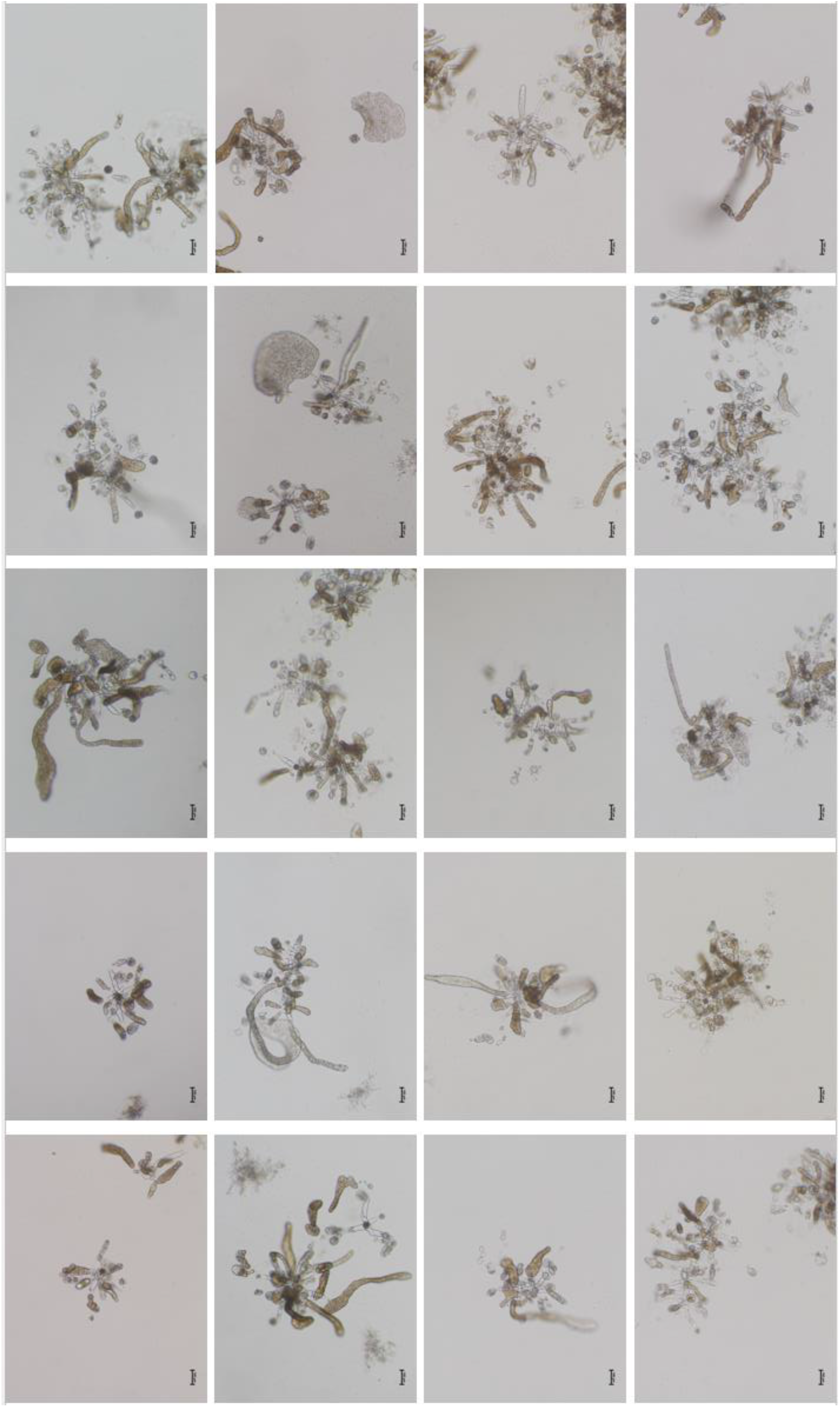
Panel with representative images of 20 independent incubations of axenic gametophytes under sporophyte growth conditions showing abnormal sporophyte development with only occasionally a long, dark sporophyte and thus healthily looking individual.

**Figure S4.**
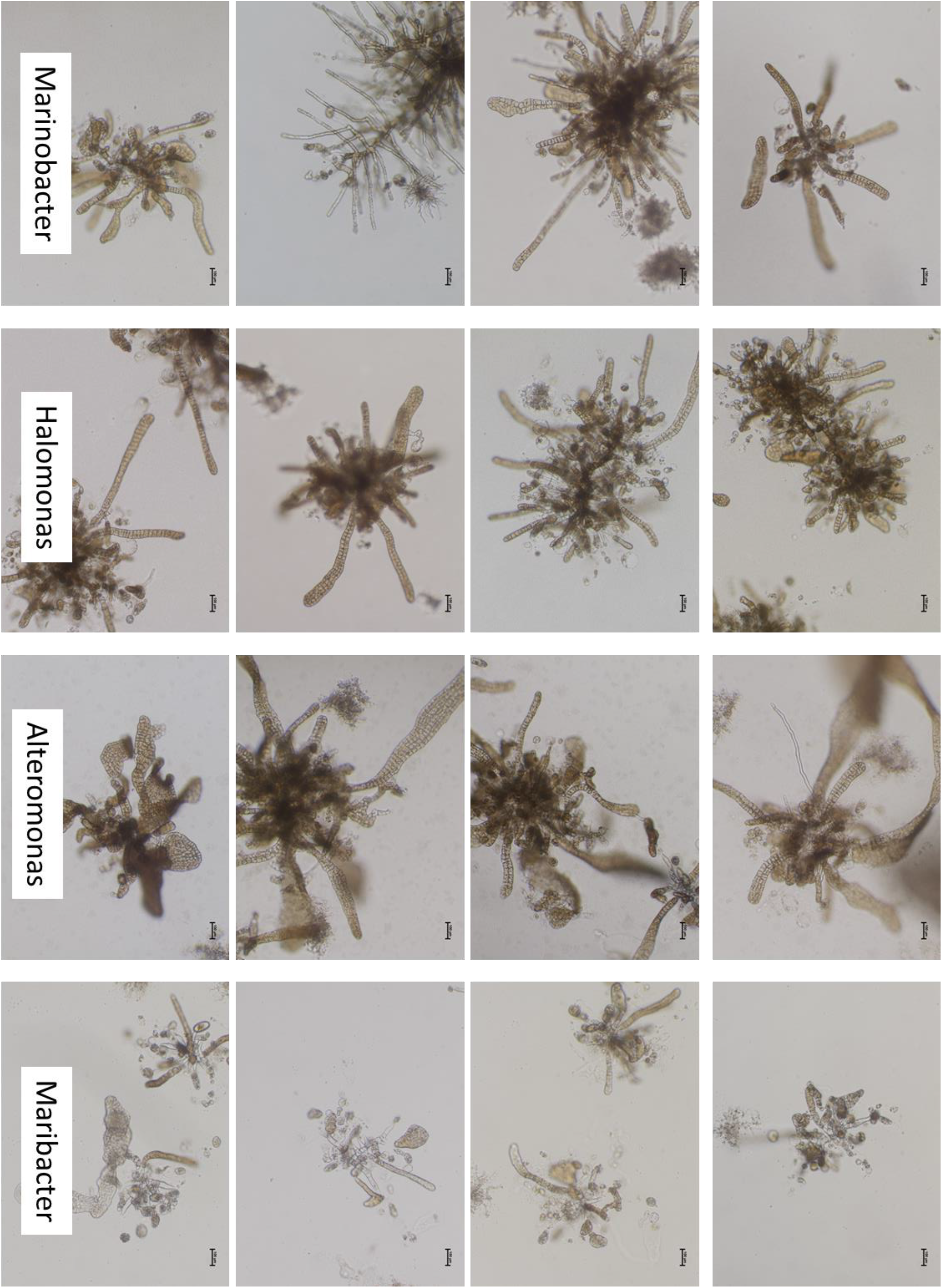
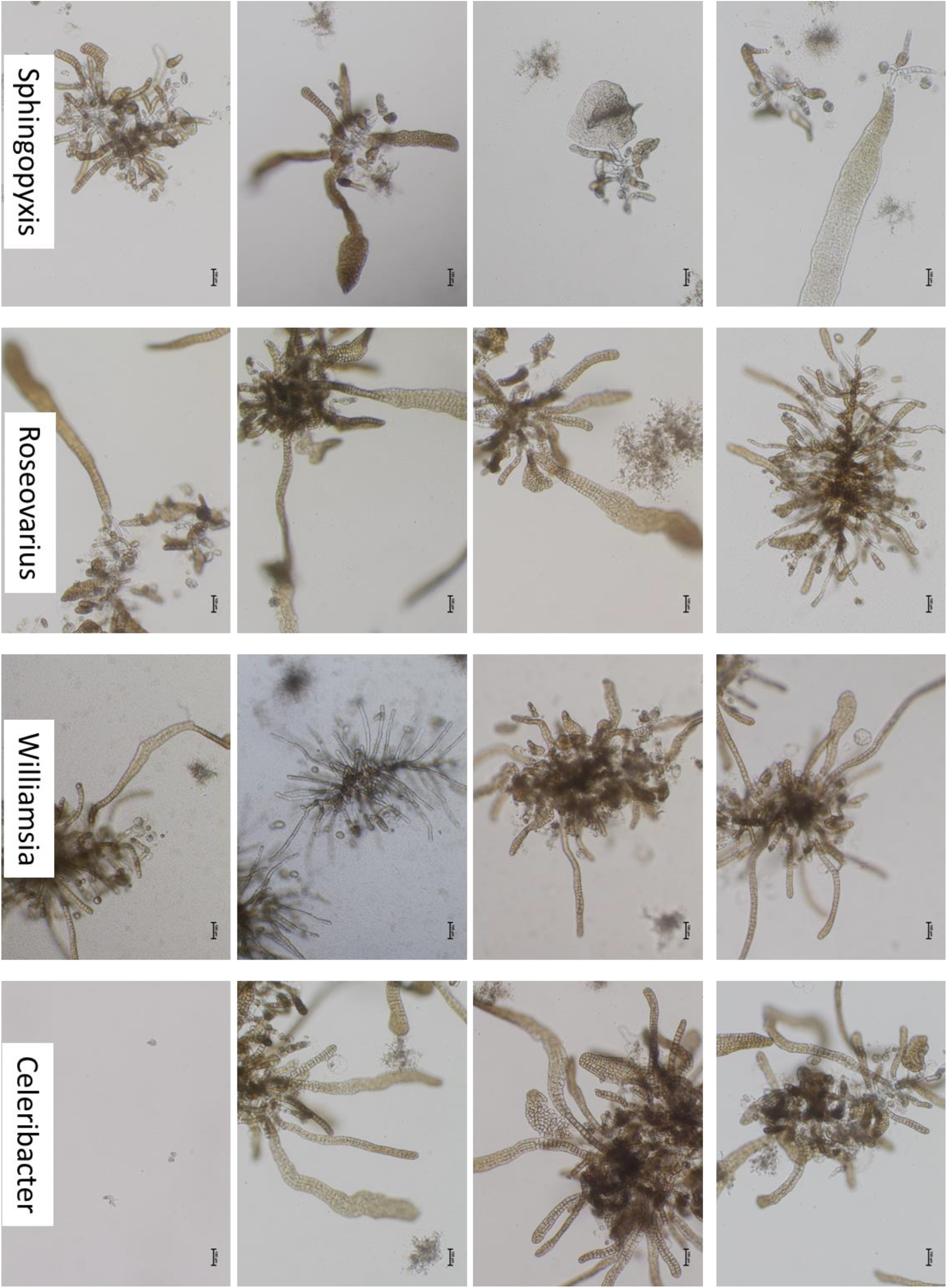
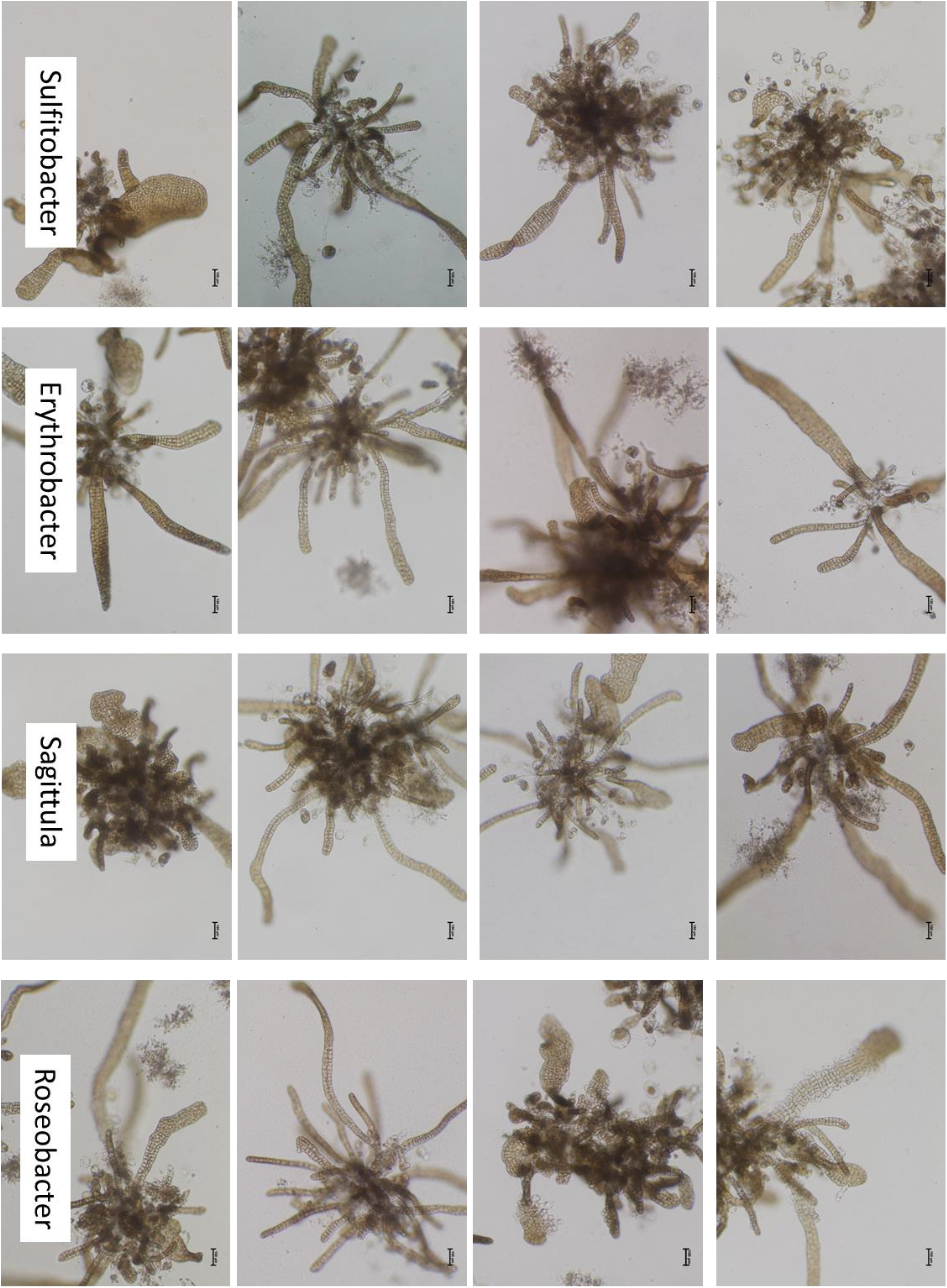
Panel with representative images of 4 independent incubations (rows from left to right) of gametophytes under sporophyte growth conditions in the presence of specific bacterial isolates. A great diversity in sporophyte number and appearance was observed, but in the presence of the majority of bacterial isolates sporophyte developed with dark pigmentation, an apical-basal axis and clearly defined cells. Note that in the presence of *P. flavimaris* and *Maribacter* sp. (MS6) sporophytes similar to axenic sporophytes (figure S4) developed and in the presence of *W*. sp. and *M. alkaliphilus*occasionally gametophytes remained in a vegetative state.

